# DNA replication stress induced by trifluridine determines tumor cell fate according to p53 status

**DOI:** 10.1101/764522

**Authors:** Yuki Kataoka, Makoto Iimori, Ryo Fujisawa, Tomomi Morikawa-Ichinose, Shinichiro Niimi, Takeshi Wakasa, Hiroshi Saeki, Eiji Oki, Daisuke Miura, Toshiki Tsurimoto, Yoshihiko Maehara, Hiroyuki Kitao

## Abstract

DNA replication stress is a predominant cause of genome instability, a driver of tumorigenesis and malignant progression. Nucleoside analog-type chemotherapeutic drugs introduce DNA damage and exacerbate DNA replication stress in tumor cells. However, the mechanisms underlying tumor cytotoxicity triggered by the drugs are not fully understood. Here, we show that the fluorinated thymidine analog trifluridine (FTD), an active component of the chemotherapeutic drug trifluridine/tipiracil, delayed DNA synthesis by human replicative DNA polymerases. FTD acted as an inefficient deoxyribonucleotide triphosphate source (FTD triphosphate) and as an obstacle base (trifluorothymine) in the template DNA strand. At the cellular level, FTD decreased thymidine triphosphate in the dNTP pool and induced FTD triphosphate accumulation, resulting in replication fork stalling caused by FTD incorporation into DNA. DNA lesions involving single-stranded DNA were generated as a result of replication fork stalling, and the p53-p21 pathway was activated. Although FTD suppressed tumor cell growth irrespective of p53 status, tumor cell fate diverged at the G2/M phase transition according to p53 status; tumor cells with wild-type p53 underwent cellular senescence via mitosis skip, whereas tumor cells that lost wild-type p53 underwent apoptotic cell death via aberrant late mitosis with severely impaired separation of sister chromatids. These results suggest that DNA replication stress induced by a nucleoside analog-type chemotherapeutic drug triggers tumor cytotoxicity by determining tumor cell fate according to p53 status.

**Significance:** This study identified a unique type of DNA replication stress induced by trifluridine, which directs tumor cell fate either toward cellular senescence or apoptotic cell death according to p53 status.

## INTRODUCTION

Accurate DNA replication is fundamental to faithful genome duplication and cellular proliferation (1). Obstacles that perturb DNA replication process induce cellular stress termed DNA replication stress (DRS). Tumor cells often lose control mechanisms of DNA replication with sustained proliferation signaling, which leads to DNA damage and chronic DRS (2). Conventional chemotherapy exerts its cytotoxic effects by introducing DNA damage and simultaneously promotes DRS by perturbing the DNA replication process (3). DRS induced by chemotherapy triggers the activation of cellular response pathways for survival in tumor cells; however, loss or suppression of the stress response can increase the susceptibility of tumor cells to catastrophic failure of proliferation. Thus, exploiting DRS is a feasible approach for cancer therapy (4,5).

DRS triggers various cellular responses in tumor cells. DRS activates ataxia telangiectasia and Rad3-related (ATR) kinase, which is recruited to replication protein A (RPA)-coated single-stranded DNA (ssDNA) at the stalled replication fork and transduces signals to numerous downstream targets (*e.g*. Chk1 is the main effector kinase) via phosphorylation to activate the S phase checkpoint (6). Thus, ATR and Chk1 preserve genome integrity by stabilizing the stalled fork and preventing origin firing (6,7). DRS also activates the Fanconi anemia (FA) pathway, which is a biochemical network that contributes to DNA repair and replication (8). FA pathway activation, which is triggered by the ATR-dependent phosphorylation of FancI, results in FancD2 monoubiquitination (9,10). In cases of moderate DRS, however, tumor cells can escape the S phase checkpoint and proceed to G2 and M phases with under-replicated DNA (UR-DNA) at hard-to-replicate DNA regions, such as common fragile sites (CFSs) (11). The UR-DNAs form ultrafine DNA bridges (UFBs) at anaphase (12) and induce genome instability phenotypes, such as lagging chromosomes and micronuclei (13). Moreover, chronic exposure to moderate DRS activates p53 and induces 4 cellular senescence-like growth arrest (14). Cellular senescence in tumor cells is also induced by anticancer chemotherapy and improves long-term outcomes (15).

Nucleoside analog-type chemotherapeutic drugs are structurally similar antimetabolites with a broad range of action, and they are clinically active in both solid tumors and hematological malignancies (16). Nucleoside analogs are efficiently incorporated into the cytoplasm of tumor cells, rapidly phosphorylated to triphosphate forms, and incorporated into DNA during normal DNA synthesis by replicative DNA polymerases. Trifluridine (FTD) is a fluorinated thymidine analog included in the clinically-approved chemotherapeutic drug called trifluridine/tipiracil (FTD/TPI; also named TAS-102) (17-20). FTD is massively incorporated into DNA without detectable DNA strand breaks, induces the phosphorylation of Chk1 at serine 345 (pS345 Chk1), a specific site phosphorylated by ATR kinase upon DRS (6), and activates the p53-p21 pathway, leading to sustained cell cycle arrest at the phase with 4N DNA content (21). However, the mechanism underlying the induction of DRS by FTD and its contribution to subsequent cell fate decisions remains to be fully elucidated.

To our knowledge, this is the first study to demonstrate that FTD, as a nucleoside analogue, delays DNA replication *in vitro*. At the cellular level, FTD stalled replication forks and subsequently generated DNA lesions including ssDNA. This led tumor cells to activate the p53-p21 pathway, skip mitosis, and fall into the persistent growth arrest characterized as cellular senescence. On the other hand, p53 knock-out tumor cells showed aberrant mitosis with severely impaired sister chromatid separation that elicited apoptotic cell death. These data indicate that DRS induced by FTD treatment triggers tumor cytotoxicity by determining tumor cell fate according to p53 status.

## MATERIALS AND METHODS

### Cell culture and reagents

HCT-116 cells were purchased from American Type Culture Collection in 2011. DLD1 cells were provided by Taiho Pharmaceutical Co. Ltd. (22). A549 cells were provided by Dr. M. Takeshita (Kyushu University) (21). All cells were authenticated by short tandem repeat analysis and confirmed negative for *Mycoplasma* infection with the MycoAlert Mycoplasma Detection Kit (Lonza). HCT-116 and A549 cells were cultured in DMEM, and DLD1 cells were cultured in RPMI1640 supplemented with 10% FBS, 100 U/mL penicillin, and 100 mg/mL streptomycin at 37°C in 5% CO_2_. The following reagents were used: RO-3306, thymidine, chlorodeoxyuridine (CldU), iodedeoxyuridine (IdU), bromodeoxyuridine (BrdU), dTTP, BrdUTP (Sigma), FTD (Tokyo Chemical Industry), and FTD-TP (Movarek).

### DNA fiber analysis

DNA fiber analysis was performed as described previously (23) with some modifications. For nascent DNA labeling, DLD1 cells were cultured in the presence of 20 μM CldU for 20 min, washed twice with fresh medium, and cultured in the presence of 20 μM IdU or 20 μM FTD for the indicated times. After double labeling, cell suspensions were spotted on slides, air-dried, and lysed with cell lysis solution (200 mM Tris-HCl, pH7.5, 50 mM EDTA, and 0.5% SDS). Following cell lysis, the slides were tilted to 15°to allow the DNA fibers to spread along the slides. After fixation with methanol/acetic acid (3:1) and DNA denaturation with 2.5 N HCl, the DNA fibers on slides were immunostained with two different anti-BrdU antibodies (Table S1), anti-mouse IgG conjugated with Alexa Fluor 568, and anti-rat IgG conjugated with Alexa Fluor 488 (Molecular Probes) at 1:400 dilution. Slides were mounted with Vectashield (H-1000; Vector lab).

### *In vitro* DNA polymerase assay

The proteins used for the *in vitro* DNA polymerase assay (Fig. S2) were purified as described previously (24-26). Human DNA polymerase δ/ε activity was measured with reference to the incorporation of [α-^32^P] dAMP. The reaction mixture (10 μL) contained 25 mM Hepes-NaOH (pH 7.8), 0.1 mg/mL BSA, 0.5 mM DTT, 10 mM Mg(CH_3_COOH)_2_, 2 mM ATP, 100 μM each of dGTP, dCTP, dATP, the indicated concentrations of dTTP or dTTP analog, 0.625–1.25 μM [α-^32^P] dATP, 12.4 fmol (90 pmol for nucleotides) of singly primed M13mp18 DNA (the 90-mer primer: 5′-AGGCGGTCAGTATTAACACCGCCTGCAACAGTGCCACGCTGAGAGCC AGCAGCAAATGAAAAATCTAAAGCATCACCTTGCTGAACCTCA-3′ is complementary to nucleotide positions 4833 and 4922), 3.5 pmol replication protein A for Polδ or 3.5 pmol ssDNA binding protein for Polε, 1 pmol proliferating cell nuclear antigen (PCNA), 60 fmol replication factor C, and the indicated amounts of polymerase. After incubation at 37°C for 30 min, reaction mixtures were immediately chilled on ice, and 7 μL samples were spotted on Whatman DE81 paper (GE Healthcare). The unincorporated nucleotides were washed four times with 0.5 M Na_2_HPO_4_, and the incorporated [α-^32^P] dAMP adsorbed onto the paper was measured by Cherenkov counting with a liquid scintillation counter (Beckman Coulter).

To measure DNA synthesis on the oligonucleotides, the 5′ end of the oligonucleotide primer was radiolabeled with [γ-^32^P] ATP, annealed with the template oligonucleotide, and subjected to *in vitro* DNA polymerase reaction at 37°C for the indicated times. The reaction was stopped by adding a bromophenol blue/xylene cyanate-formamide EDTA solution. After boiling at 95°C for 3 min, the samples were loaded onto 15% acrylamide gels containing 7% urea and electrophoresed at 30W for 60 min. The gels were fixed with 15% methanol/15% acetate solution for 10 min, rinsed with tap water, and dried on 3MM paper. Radioactivity was detected using BAS2000 (GE Healthcare).

### Quantification of dTTP, FTD-TP, and BrdUTP using liquid chromatography/triple-stage quadrupole mass spectrometry (LC-QqQ-MS)

Intracellular dTTP, FTD-TP, and BrdUTP were quantified using LC-QqQ-MS as described previously (27). FTD-TP and BrdUTP were detected with optimized selective reaction-monitoring transitions in negative ionization mode as follows: FTD-TP: precursor ion [*m/z*]/product ion [*m/z*] = 535/159, 535/79, and 535/257 and BrdUTP: precursor ion [*m/z*]/product ion [*m/z*] = 546.5/159.

### Generation of *TP53*-deficient HCT-116 cells

First, a 20-mer sgRNA target sequence (5′-CTCAGAGGGGGCTCGACGCT-3′) at exon 2 of the *TP53* gene was designed (28) and cloned into pX330 (Addgene #42230), which was a gift from Dr. Feng Zhang (29). The donor DNA plasmid was constructed from the PCR fragment [∼1,300-bp of the *TP53* genomic region including the sgRNA target sequence amplified by PCR (forward: 5′-ACTATATCCTTGTTAACAGGAGGTGGGAGC-3′; reverse: 5′-AAGGGTGAAGAGGAATCCCAAAGTTCCAAAC-3′)] in pCR4-TOPO [Thermo Fisher Scientific (hereafter, Thermo)], and its 130-bp *Bam* HI fragment that includes the sgRNA target sequence was replaced with the 2,500-bp *Bam* HI fragment containing a puromycin-resistant gene cassette (30). The above two plasmids were co-transfected into HCT-116 cells using 4D Nucleofector (Lonza). The puromycin-resistant clones were screened by genomic PCR and sequencing.

### Generation of fluorescent ubiquitination-based cell cycle indicator (Fucci) expressing cells by lentiviral infection

cDNA encoding mKO2-hCdt1 (aa 30–120) or mAG1-hGeminin (aa 1–110) amplified from the pFucci-G1 Red plasmid (AM-V9003) or pFucci-S/G2/M Green plasmid (AM-V9016) (MBL), respectively, was cloned into the pENTR D-TOPO vector (Thermo). Each plasmid 8 was mixed with pLenti6.4/R4R2/V5-DEST and pENTR 5′/EF1αP and recombined using LR Clonase II Plus enzyme (Thermo). The lentiviruses were produced using the ViraPower Lentiviral Expression System (Thermo). To establish HCT-116-Fucci or HCT-116 *p53*^*-/-*^-Fucci cells, cells were infected with both lentiviruses encoding mKO2-hCdt1 (30–120) and mAG1-hGeminin (1–110) at a multiplicity of infection of 1 each. The infected cells were first selected by blasticidin (Thermo) treatment. The surviving cells were sorted in two steps using BD FACSAria SORP (BD Bioscience) as follows: cells emitting red fluorescence were sorted and cultured for several days, and then cells emitting green fluorescence were sorted.

### Western blot analysis

Western blot analysis was performed as described previously (21). The antibodies used are listed in Table S1. Chemiluminescence was detected using LAS 4000 mini (GE Healthcare).

### Immunofluorescence

Cells were rinsed in PBS at 37°C, fixed in 4% paraformaldehyde (PFA) for 15 min at 37°C (mitosis)(31) or 3% PFA, 2% sucrose, and 0.5% Triton X-100 for 30 min on ice (nuclear foci)(32), permeabilized in PBS containing 0.1% Triton X-100 for 5 min at 37°C, blocked in PBS containing 2% BSA for 30 min at room temperature, and incubated for 16 h at 4°C with the antibodies listed in Table S1. Secondary antibodies conjugated to Alexa Fluor 488, 568, or 647 (Molecular Probes) were used at 1:2,000 dilution. After washing in PBS containing DAPI for 5 min, the coverslips were mounted in ProLong Diamond or ProLong Glass (Thermo).

### Image acquisition

For fixed-cell experiments, fluorescence image acquisitions were performed using a Nikon A1R confocal imaging system or N-SIM super resolution imaging system controlled by NIS Elements software (Nikon). The objective lens was an oil immersion CFI SR ApoTIRF 100× 9 NA 1.49 lens, an oil immersion Plan-Apo 100× NA 1.45 lens, or a Plan-Apo 40× NA 0.95 lens (Nikon). Images were acquired as Z-stacks at 0.2-μm or 0.12-μm intervals with a confocal or a super-resolution microscope, respectively, and maximum-intensity projections were generated using NIS Elements software (Nikon).

For live-cell imaging of cell cycle progression, HCT-116-Fucci or HCT-116 *p53*^*-/-*^-Fucci cells were imaged in a chambered coverglass (Matsunami) containing phenol red-free DMEM (Gibco). Live-cell imaging was performed as described previously (31). The duration of the cell cycle phases was calculated manually.

For the quantification of nuclear foci of FancD2 and RPA32, fluorescence images were acquired using BZ-X800 (Keyence) with Plan-Apo 40×NA, and nuclear foci were counted using Hybrid Cell Count Software (Keyence).

### Animals and evaluation of antitumor activity *in vivo*

All animal studies were performed according to the guidelines and with the approval of the institutional Animal Care and Use Committee of Taiho Pharmaceutical Co., Ltd. Ethical approval (5 Mar 2019) was obtained prior to conducting the animal experiments. Male nude mice (CLEA Japan) were housed under specific pathogen-free conditions, with food and water provided ad libitum. The animals were quarantined for 1 week and then subcutaneously implanted with 1 × 10^7^ HCT-116 or HCT-116 *p53*^*-/-*^ cells on day 0. The mice were randomized on day 3 based on body weight and each treatment group consisted of six mice. FTD/TPI was prepared by mixing FTD and TPI at a molar ratio of 1:0.5 in 0.5% HPMC solution. FTD/TPI (FTD: 200 mg/kg/day) was administered orally twice daily from days 3–7, days 10–14, and days 17–21 at approximately 6-hour intervals. For the control group, vehicle (0.5% HPMC solution) was administered at 10 mL/kg.

### Statistical analysis

The statistical analysis was performed using GraphPad Prism (GraphPad Software) or EXSUS (CAC Croit Corp.) software. The Mann-Whitney *U* test was used in Figs. 1B and 5C, the Kruskal-Wallis test in Fig. 3D and E, and the unpaired *t*-test in Figs. 4F, 4G, 6B, and 6D.

**Fig. 1.**
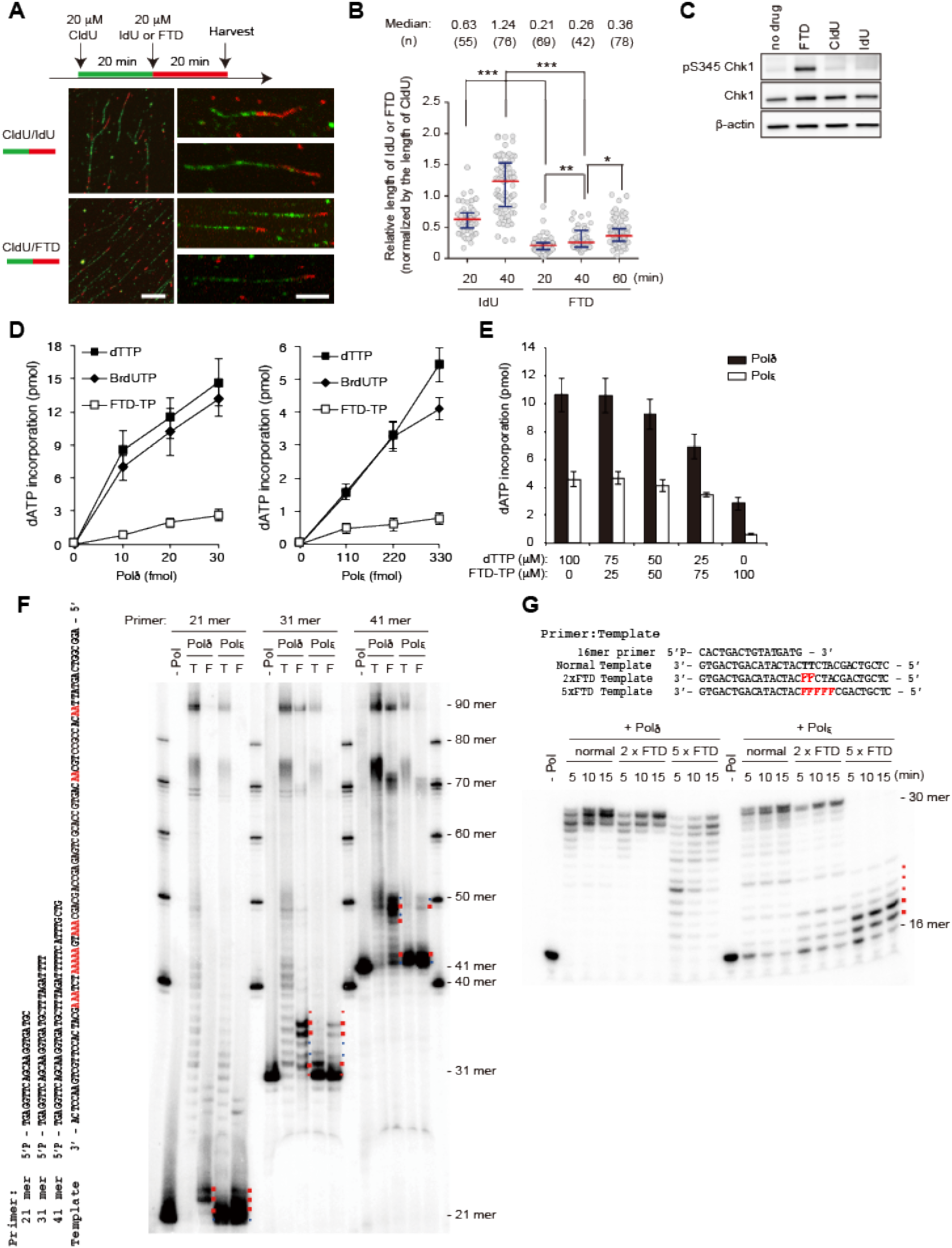
FTD retards replication fork progression. **A**, DNA fiber analysis; DLD1 cells were incubated with 20 μM CldU for 20 min and either with 20 μM IdU (upper) or 20 μM FTD (lower) for 20 min. Representative images are enlarged. Scale bars indicate 10 μm (left) and 5 μm (enlarged). **B**, Scatter plot of IdU-/CldU-tract length ratios for individual replication forks in **A**. Red lines denote median and blue whiskers extend to the quartiles. Mann-Whitney *U-*test. *: *p<*0.05, **: *p*<0.01, ***: *p*<0.001. **C**, Western blot; DLD1 cells were incubated in the presence of 20 μM nucleoside analogs for 1 hour. **D**, The *in vitro* DNA synthesis rate of replicative polymerases Polδ (left) and Polε (right), in the presence of dTTP, BrdUTP or FTD-TP. DNA synthesis was measured by [α-^32^P] dATP incorporation into the nascent strand DNA using M13mp18 ssDNA plasmid as a template. Error bars represent standard deviation (SD) of three independent experiments. **E**, Relative *in vitro* DNA synthesis rates in the presence of the dTTP/FTD-TP mixture. Error bars represent SD of three independent experiments. **F**, The *in vitro* DNA synthesis of Polδ and Polε in the presence of FTD-TP. Sequences of 5′-radiolabeled primers (21-mer, 31-mer or 41-mer) and template 90-mer oligonucleotides of the *in vitro* DNA synthesis analysis are shown on the left. Repetitive adenine sequences in the template oligonucleotide are marked by red. DNA synthesis from radiolabeled primers of the 90-mer oligonucleotide template is shown on the right. The DNA synthesis reaction catalyzed by Polδ and Polε was performed in the presence of dTTP (T) or FTD-TP (F). The bands representing DNA synthesis stopped at repetitive adenine sequences are marked by red dots. **G**, The *in vitro* DNA synthesis catalyzed by Polδ and Polε from a DNA template containing FTD. Sequences of the 5′-radiolabeled 16-mer primer and template 31-mer oligonucleotides (normal, 2×FTD, 5×FTD template) of the *in vitro* DNA synthesis analysis are shown on the upper and repetitive FTD (F) sequences in the template oligonucleotide are marked by red. DNA synthesis from radiolabeled primers of the 31-mer oligonucleotide template is shown on the lower. The DNA synthesis reaction catalyzed by Polδ and Polε was performed for the indicated times (min). Reaction in the absence of replicative polymerase is shown by –Pol.

**Fig. 2.**
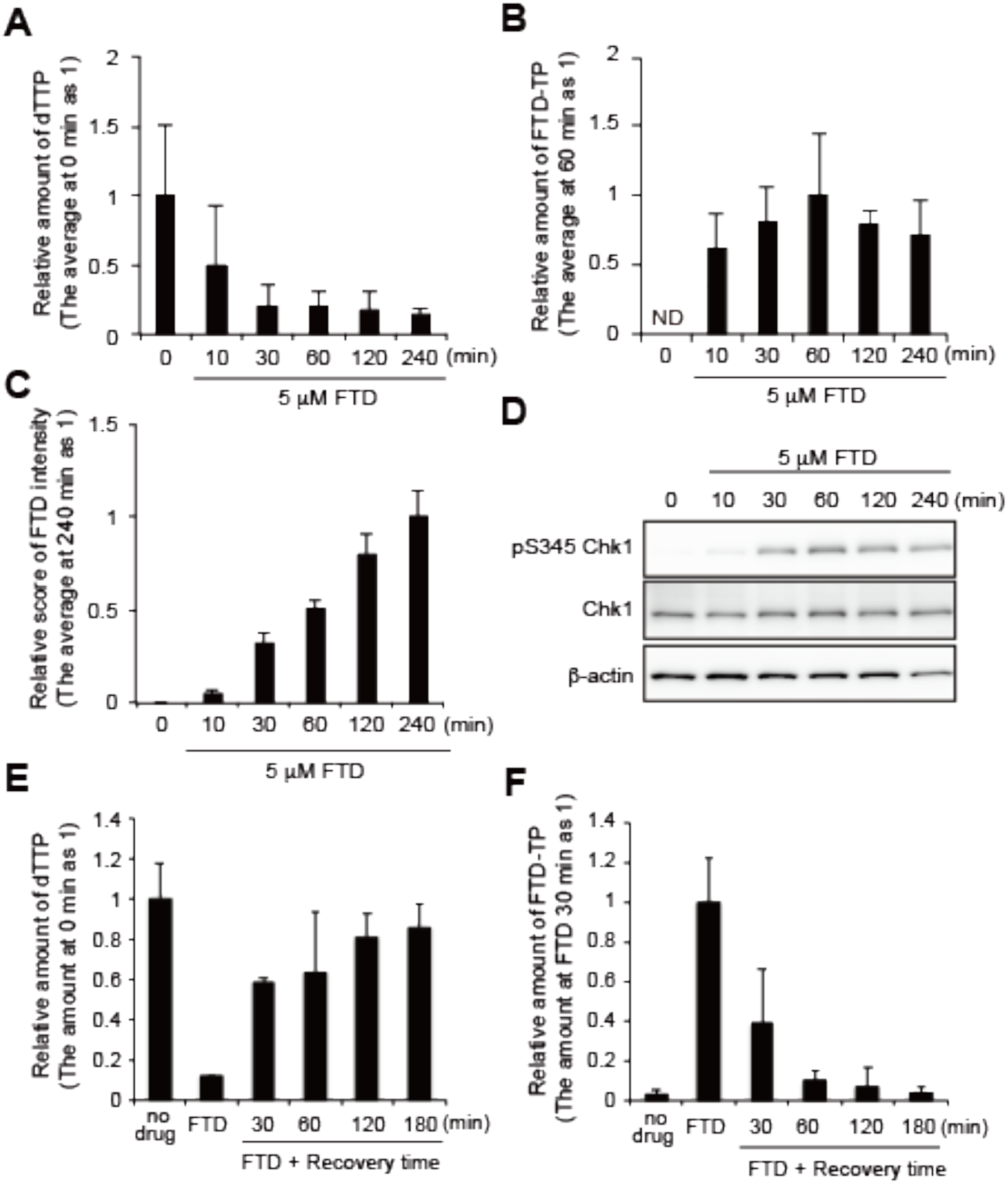
Effect of FTD on the cellular and metabolomic states. **A, B**, Relative amounts of cellular dTTP (**A**) and FTD-TP (**B**). The relative scores were calculated by considering the average amount at 0 min (**A**) or 60 min (**B**) as 1. ND: not detectable. **C**, FTD incorporation into DNA. The relative scores were calculated by considering the average amount at 240 min as 1. **D**, Western blot; HCT-116 cells were treated with 5 μM FTD and harvested at the indicated time points. pS345 Chk1: Chk1 phosphorylation at Ser345. **E, F**, Relative amounts of cellular dTTP (**E**) and FTD-TP (**F**) after the change to drug-free media. The relative scores were calculated by considering the average amount at 0 min (**E**) or 5 μM FTD 30 min (**F**) as 1. Error bars represent SD of three independent experiments.

**Fig. 3.**
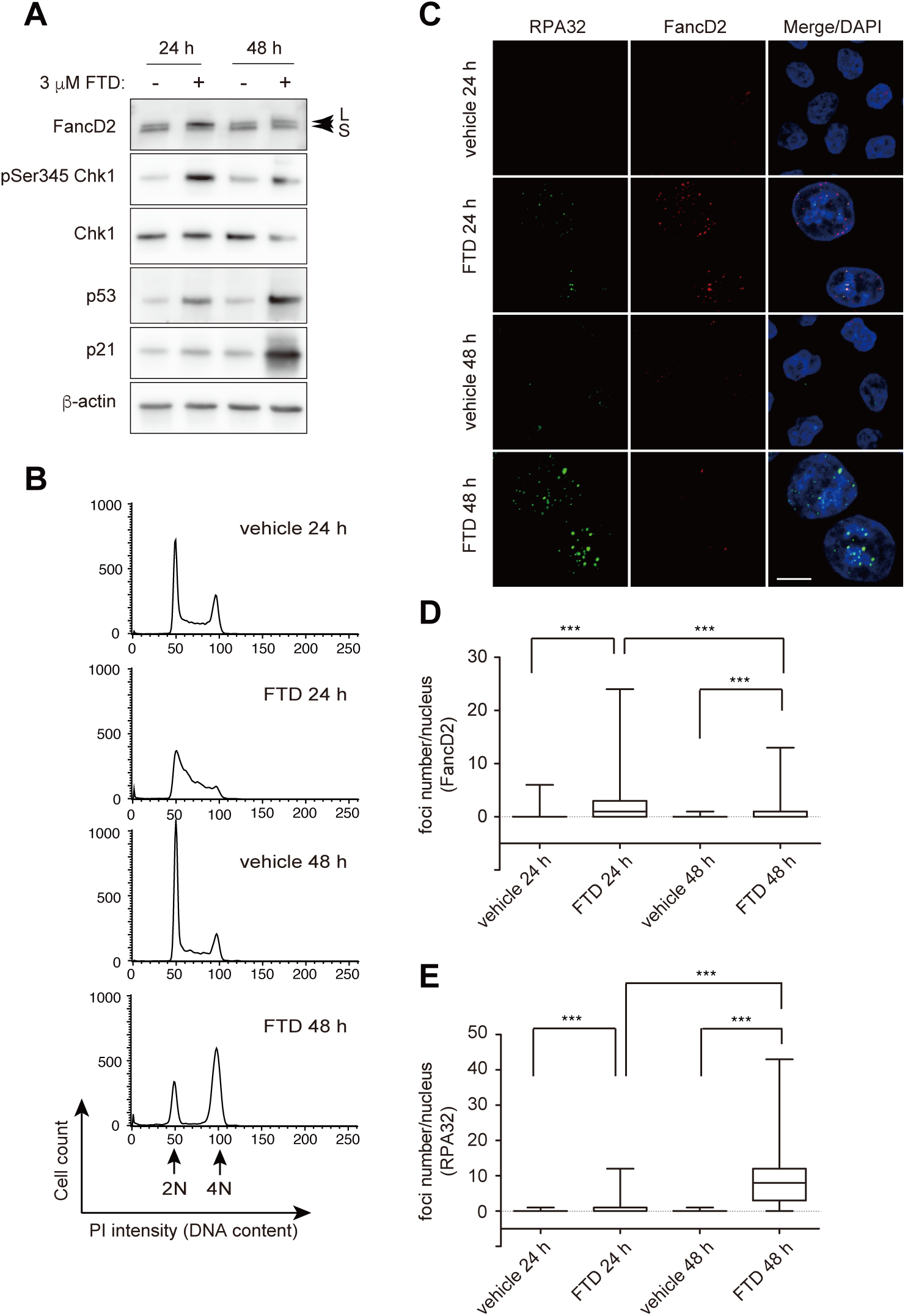
DNA damage response to FTD in HCT-116 cells. **A**, Western blot; HCT-116 cells were treated with 3 μM FTD and harvested at the indicated time points. L and S indicate large (monoubiniquitinated) and small (non-ubiquitinated) isoforms of FancD2. **B**, histogram; the position of cells with 2N and 4N DNA content are indicated by arrows. **C**, immunofluorescence images of FancD2 and RPA32; scale bar, 10 μm. **D, E**, Box plot; the number of FancD2 (**D**) and RPA32 (**E**) foci per each nucleus are plotted. The box ends are the quartiles, the horizontal lines inside the box are the median, and the whiskers extend out to the farthest points. Kruskal-Wallis test. ***: *p*<0.001.

**Fig. 4.**
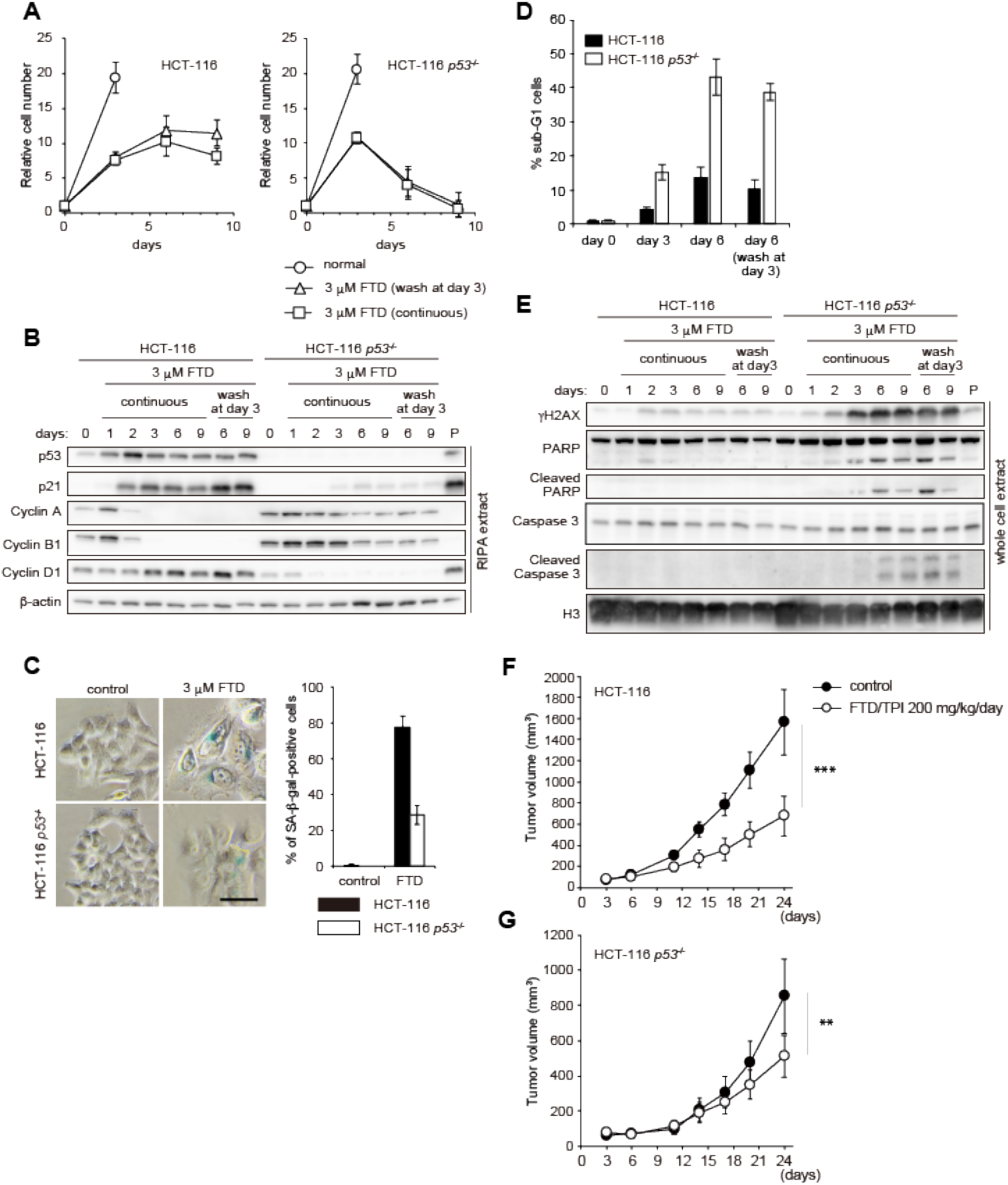
Outcomes of HCT-116 and HCT-116 *p53*^*-/-*^ cells exposed to FTD. **A**, Growth curve; Cell growth was measured by crystal violet staining on the indicated days. Representative images are shown in Fig. S6. Error bars represent SD of three independent experiments. **B**, Western blot analysis of RIPA extracts; HCT-116 and HCT-116 *p53*^-/-^ cells were treated with 3 μM FTD either continuously (continuous) or for 3 days (wash at day 3) for the indicated days and harvested. P represents a positive control (HCT-116 3 μM FTD for 3 days). **C**, Senescence associated β-galactosidase (SA-β-gal) activity of HCT-116 and HCT-116 *p53*^-/-^ cells on day 3; The black bar represents 50 μm. The graph shows the percentage of SA-β-gal-positive cells. Error bars represent the SD of three independent experiments. **D**, Western blot analysis of whole cell extracts; HCT-116 and HCT-116 *p53*^-/-^ cells were treated with 3 μM FTD as shown in (**B**). **E**, Sub-G1 population. HCT-116 and HCT-116 *p53*^-/-^ cells were treated with 3 μM FTD and harvested on the indicated days. Ethanol-fixed samples were stained with propidium iodide and the percentage of sub-G1 cells was measured. Error bars represent the SD of three independent experiments. **F**,**G**, Growth curve of HCT-116 (**F**) and HCT-116 *p53*^-/-^ (**G**) xenografts. Error bars represent the SD of 6 individual mice. Statistical analysis was done at day 24. Unpaired *t*-test. **: *p*<0.01, ***: *p*<0.001.

**Fig. 5.**
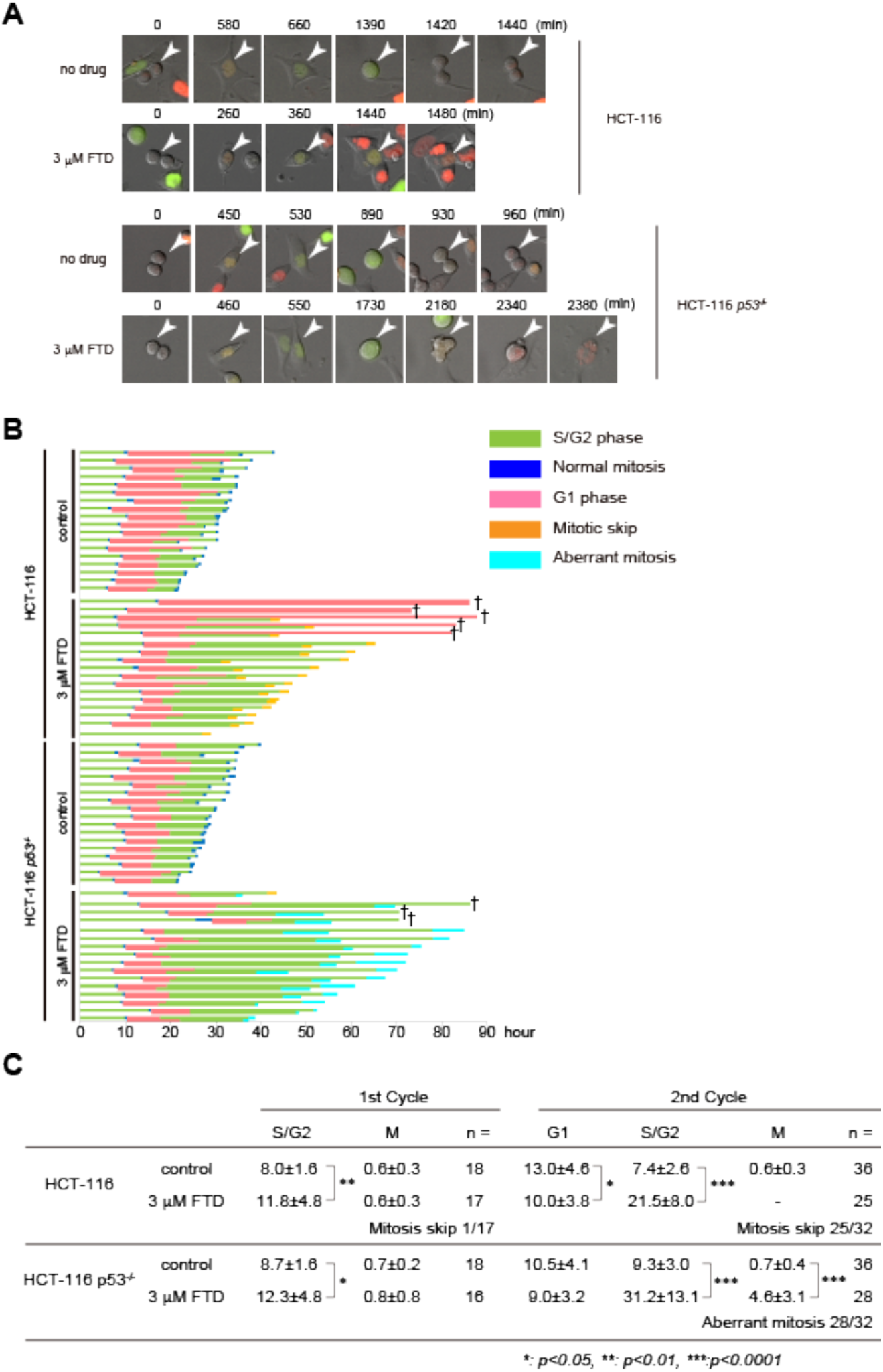
Single-cell live imaging of Fucci cells in response to FTD. **A**, Representative images of HCT-116-Fucci and HCT-116 *p53*^-/-^-Fucci cells in the presence or absence of 3 μM FTD. Arrowheads indicate identical cells at each time point. **B**, Cell cycle phase and duration in HCT-116-Fucci and HCT-116 *p53*^-/-^-Fucci cells in the presence or absence of 3 μM FTD. Each bar represents one cell that was in G1 phase (red color) at the start point. †: analysis terminated. **C**, Duration of each cell cycle phase. Mean hours ± SDs are shown. Mann-Whitney *U-*test. *: *p*<0.05, **: *p*<0.01, ***: *p*<0.001.

**Fig. 6.**
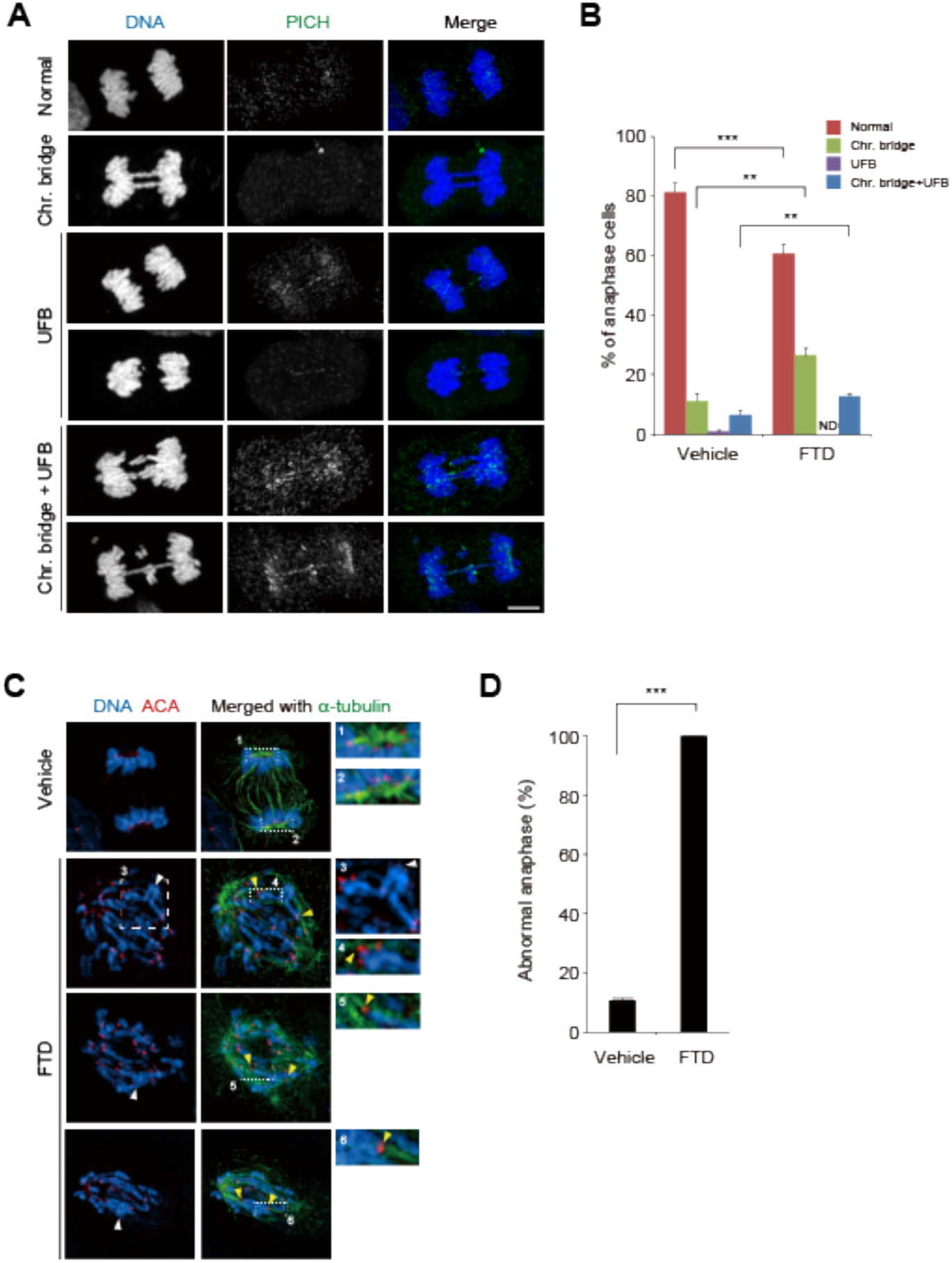
Aberrant chromosomal structures of HCT-116 *p53*^*-/-*^ cells in response to FTD. **A**, Representative immunofluorescence images of anaphase HCT-116 *p53*^-/-^ cells, which were either untreated (Normal, and UFB) or cultured in the presence of 3 μM FTD and 9 μM RO-3306 for 16 hours and released into fresh media for 50 min (Chr. Bridge, and Chr. Bridge+UFB). **B**, Quantitative data of **A**. Error bars represent SD of three independent experiments. Unpaired *t-*test. **: *p*<0.01, ***: *p*<0.001. **C**, Representative super-resolution immunofluorescence images of anaphase HCT-116 *p53*^-/-^ cells, which were cultured in the presence of 3 μM FTD for 60 h. The centromeres (ACA) captured by spindles are indicated by yellow arrowheads. Sister chromatid pairs with entangled chromosome arms are indicated by white arrowheads. Enlarged images of rectangles with dashed lines are shown in the inlets. **D**, Quantitative data of abnormal anaphase. Error bars represent SD of three independent experiments. Unpaired *t-*test. ***: *p*<0.001.

## RESULTS

### FTD stalls replication forks and activates the DNA damage response

In a previous study, we showed that FTD induces pS345 Chk1, an indicator of the DNA damage response to DRS, during its misincorporation into DNA (21). We hypothesized that the process of FTD misincorporation itself would cause DRS, and FTD-containing replication fork progression would be retarded. To evaluate the speed of individual active replication forks, we performed DNA fiber analysis (23). Active elongating replication forks were labeled with CldU, and their elongation was measured by subsequent labeling with IdU or FTD (Fig. 1A and B) because IdU and FTD are recognized by B44 but not by BU1/75 (Fig. S1A and B) (33). FTD-containing forks were significantly shorter than IdU-containing replication forks (Fig. 1A and B). Furthermore, FTD, but not CldU or IdU, induced pS345 Chk1 (Fig. 1C). These results indicate that FTD stalled replication forks during its incorporation into DNA and activated the DNA damage response.

### FTD impedes replicative DNA polymerases *in vitro*

We hypothesized that FTD triphosphate (FTD-TP) would be incorporated into DNA during replication catalyzed by replicative DNA polymerases (Polδ and Polε) with low efficiency, which could slow the elongation of the nascent DNA strand. To test this possibility, DNA synthesis was evaluated by measuring the incorporation of radioactive [α-^32^P] dATP in an *in vitro* reconstituted DNA replication assay using purified proteins (Fig. S2A–2C). The [α-^32^P] dATP incorporation rate was significantly lower in the presence of FTD-TP than in the presence of dTTP or BrdUTP (Fig. 1D). In addition, the low [α-^32^P] dATP incorporation rate in the presence of FTD-TP was almost completely rescued when 50% of FTD-TP was replaced by dTTP (Fig. 1E). These results indicate that the incorporation rate of FTD-TP into the nascent DNA strand was less efficient than that of dTTP or BrdUTP.

To identify the specific sequence in the template DNA strand at which the synthesis of the nascent DNA strand is impeded in the presence of FTD-TP, we performed an *in vitro* DNA replication assay with a defined DNA template. Synthesis of the nascent DNA strand in the presence of FTD-TP was strongly and specifically impeded at positions of repetitive adenine sequences (Fig. 1F), whereas synthesis was only slightly impeded at positions of dispersed adenine sequences (Fig. S3).

We further examined whether the elongation of the nascent DNA strand was retarded when Polδ and Polε encounter FTD in the template DNA strand. Elongation was significantly suppressed when either Polδ or Polε was used to replicate a template DNA strand containing five repetitive FTDs at the 3′ vicinity of the primer end (Fig. 1G). Collectively, these results indicate that FTD induced DRS by impeding the progression of replicative DNA polymerases during its incorporation into the nascent strand DNA as well as when FTD was present in the template DNA strand.

### FTD addition rapidly decreases dTTP and increases FTD-TP in the cellular dNTP pool

Next, we examined whether FTD-TP was generated when cells were cultured in the presence of FTD. In HCT-116 cells cultured in the presence of various concentrations of FTD for 60 min, FTD incorporation was detected (Fig. S4A), and dTTP decreased and FTD triphosphate (FTD-TP) increased in a concentration-dependent manner (Fig. S4B and C). In HCT-116 cells cultured in the presence of FTD, dTTP decreased rapidly (in 30 min) as FTD-TP increased in the cellular dNTP pool (Fig. 2A and B). At this time point, FTD was being continuously incorporated into DNA (Fig. 2C) and induced pS345 Chk1 (Fig. 2D). When FTD was removed from the medium, dTTP recovered rapidly as FTD-TP disappeared from the dNTP pool (Fig. 2E and F). These data indicate that FTD-TP was generated when cells were cultured in the presence of FTD and that it activated the DNA damage response during its incorporation into DNA.

### DNA lesions including ssDNA are generated as a result of FTD-induced replication fork stalling

Next, we monitored cellular responses in HCT-116 cells cultured in the presence of FTD. At 24 h, FTD induced pS345 Chk1 and FancD2 monoubiquitination (Fig. 3A), indicating activation of ATR kinase and the Fanconi anemia DNA repair pathway in response to stalled replication forks. Consistently, major populations of cells were in S phase at 24 h (Fig. 3B). At 48 h, when most cells showed 4N DNA content (Fig. 3B), however, pS345 Chk1 and FancD2 monoubiquitination had decreased and p53 and p21 had increased (Fig. 3A). This result suggests that FTD transiently stalled replication forks and activated the ATR-dependent DNA damage response, which in turn triggered activation of the p53-p21 pathway probably via DNA lesions arising from improperly processed stalled replication forks.

Failure to complete DNA replication during S phase produces ssDNA-containing UR-DNA, which leads to DNA lesions that are transmitted across cell generations if they are not repaired properly (34-36). To investigate whether FTD generated ssDNA, we co-immunostained the ssDNA binding protein RPA32 and FancD2 in FTD-treated HCT-116 cells (Fig. 3C). We observed a modest increase of RPA32 nuclear foci at 24 h, when pan-nuclear FancD2 foci were formed. At 48 h, however, when only a small number of FancD2 nuclear foci were observed, the number and intensity of RPA32 nuclear foci increased dramatically (Fig. 3D and E). These data suggest that FTD generated ssDNA that persisted because of replication fork stalling.

### FTD induces senescence in tumor cells with wild-type p53 and apoptosis in p53 knockout tumor cells

We previously showed that FTD induces p53-dependent sustained cell cycle arrest at the phase with 4N DNA content (21). To elucidate the role of p53 in the FTD-induced DNA 14 damage response, we generated isogenic *TP53* gene knock-out HCT-116 cell lines using the CRISPR/Cas9 system (Fig. S5). In the presence of FTD, cell proliferation was similarly suppressed in HCT-116 and HCT-116 *p53*^-/-^ cells until day 3. In HCT-116 cells, proliferation ceased completely on day 6, and cell number did not change until day 9. However, in HCT-116 *p53*^-/-^ cells, cell number started to decrease after day 3 and most cells had disappeared by day 9 (Figs. 4A and S6). The same results were obtained when FTD was removed from the cell culture media of both cell lines on day 3 (Fig. 4A), indicating that cell fate was determined before day 3.

We next examined the cellular response to FTD in HCT-116 and HCT-116 *p53*^-/-^ cells (Fig. 4B). HCT-116 cells showed accumulation of p53 and p21 and almost undetectable expression of cyclin B1 and cyclin A proteins on day 3 at the time when accumulation of cyclin D1 was observed (Fig. 4B), suggesting that a significant proportion of cells were in G1 phase. In HCT-116 *p53*^-/-^ cells, however, p21 and cyclin D1 did not accumulate, and cyclin B1 and cyclin A levels did not decrease (Fig. 4B). Furthermore, similar results were obtained using A549 (p53 wild type) and DLD1 (p53 mutant) cells (Fig. S7A–C). These data indicate that FTD activated the p53-p21 pathway and suppressed the growth of cells expressing wild-type p53, resulting in the accumulation of cells in G1 phase, whereas in cells without p53, FTD caused tumor cell death.

Chronic DRS activates p53 and induces cellular senescence-like growth arrest (14). Consistently, on day 3 of FTD treatment, most HCT-116 cells were senescence-associated β-galactosidase (SA-β-gal)-positive, whereas only a limited population of HCT-116 *p53*^-/-^ cells showed this phenotype (Fig. 4C). Similar results were obtained with A549 and DLD1 cells (Fig. S7D). By contrast, on day 6 of FTD treatment, the number of cells in the sub-G1 population was greater in HCT-116 *p53*^-/-^ cells than in HCT-116 cells (Fig. 4D). Immunoblot analysis showed a dramatic upregulation of markers of DNA damage (γH2AX) and apoptosis 15 (cleaved PARP and caspase 3) in HCT-116 *p53*^-/-^ cells, but not in HCT-116 cells (Fig. 4E). These data indicate that cellular senescence and apoptotic cell death were predominantly induced in FTD-treated HCT-116 and HCT-116 *p53*^-/-^ cells, respectively.

Lastly, we examined the effect of tumor p53 status on the response to FTD/TPI treatment in an *in vivo* xenograft mouse model. FTD/TPI treatment significantly suppressed the growth of HCT-116 and HCT-116 *p53*^*-/-*^ xenograft tumors (Fig. 4F and G), indicating that FTD/TPI exerts its tumor suppressive effect irrespective of p53 status.

### Cell fate is determined according to p53 status at the G2/M transition following an FTD-induced extended S/G2 phase

We showed that FTD treatment determined cell fate in a p53-dependent manner. To explore this phenomenon in detail, time-lapse analysis of HCT-116 and HCT-116 *p53*^*-/-*^ cells was performed using the Fucci system to visualize cell cycle status (37). DNA damage-induced cellular senescence-like G1 phase arrest proceeds via mitosis skip, which is a G2-to-G1 phase transition without mitotic cell division (38). Consistent with this finding and the results of the SA-β-gal assay in HCT116 cells (Fig. 4C), most HCT116-Fucci cells showed a mitosis skip phenotype, whereas a minor proportion of HCT116-Fucci cells displayed G1 phase arrest after normal mitosis in response to FTD treatment (Fig. 5A and B). Furthermore, although HCT116 *p53*^*-/-*^ cells entered mitosis, the duration of mitosis was severely extended, and cells exited mitosis and entered the next G1 phase without separation into daughter cells (Fig. 5A–C). Intriguingly, FTD treatment markedly extended the duration of the second S/G2 phase rather than the first S/G2 phase in both HCT-116 and HCT-116 *p53*^*-/-*^ cells (Fig. 5B and C). Collectively, these data indicate that FTD treatment specifically slows the progression of S/G2 phase irrespective of p53 status, although p53 is specifically involved in cell fate determination at the G2/M phase transition after the second G2 phase.

### Severe chromosomal bridges are induced by FTD during late mitosis in p53-null cells

Upon mild perturbation of DNA replication, sister chromatids are frequently interlinked at fragile sites, which are located at genetic loci with intrinsic replication difficulties, by BLM/PICH-associated UFBs (12). We thus investigated whether FTD affects the formation of sister chromatid interlinks. First, to evaluate the effect of FTD treatment at the first S/G2 phase on sister chromatid separation in the following anaphase, HCT-116 *p53*^*-/-*^ cells were exposed to FTD in the presence of RO-3306, a Cdk1 inhibitor that perturbs the G2 to M phase transition (39), and then released into mitosis in fresh medium for 50 min to enrich anaphase cells (Fig. 6A). Most anaphase cells showed separation of sister chromatids to the spindle poles; however, the appearance of PICH-associated UFBs, most of which also contained chromosomal bridges, was also significantly increased (Fig. 6B). These results indicate that FTD treatment-induced DRS during FTD-TP incorporation into nascent DNA strands partially disturbed the separation of several individual sister chromatids, it was not strong enough to prevent the process of chromosome segregation.

Next, we observed anaphase chromosomes during the second mitosis in the presence of FTD, most of which incorporated FTD into the template DNA strand. HCT-116 *p53*^*-/-*^ cells were exposed to FTD for 60 h, and chromosomal structures were observed in anaphase. To differentiate anaphase cells with unseparated sister chromatids from prometaphase cells, cells were immunostained for cyclin B1, and only cyclin B1-negative cells that had already proceeded into anaphase were evaluated (Fig. S8). Most anaphase cells showed no separation of sister chromatids to the spindle poles (Fig. 6C and D). In addition, super-resolution microscopy of centromere and microtubule staining showed that each kinetochore was captured by mitotic spindles (Fig. 6C, insets 4–6), similar to kinetochores with sister chromatids separated to the spindle poles as observed in normal anaphase cells (Fig. 6C, insets 1 and 2). Furthermore, closer observation of each chromosome revealed that a 17 significant population of sister chromatid pairs showed interlinking along chromosomal arms (Fig. 6C, inset 3), whereas the corresponding sister kinetochores were captured by spindles from the opposite spindle poles and were already separated (Fig. 6C, inset 4). These data indicate that, upon severe DRS induced by FTD treatment during the second S phase, p53 knock-out cells displayed severe defects in sister chromatid separation at anaphase, but not in the capture of kinetochores by spindles.

## DISCUSSION

In this study, we elucidated the mechanism underlying tumor cytotoxicity induced by the fluorinated thymidine analog-type chemotherapeutic drug, FTD. FTD treatment replaced dTTP in the dNTP pool with FTD-TP, which slowed DNA synthesis by replicative DNA polymerases. Thus, FTD slowed DNA replication fork progression and subsequently generated DNA lesions, including ssDNA, and activated p53, thereby inducing cellular senescence. In the absence of p53, FTD triggered apoptotic cell death by inducing aberrant mitosis associated with severely unseparated sister chromatids. Because tumor cells show high levels of DRS and depend on their response to DRS, exploiting DRS is a feasible approach for cancer therapy (3-5). As a chemotherapeutic drug, FTD is ideal because it exacerbates cellular DRS to exert cytotoxic effects irrespective of p53 status (Fig. 7).

**Fig. 7.**
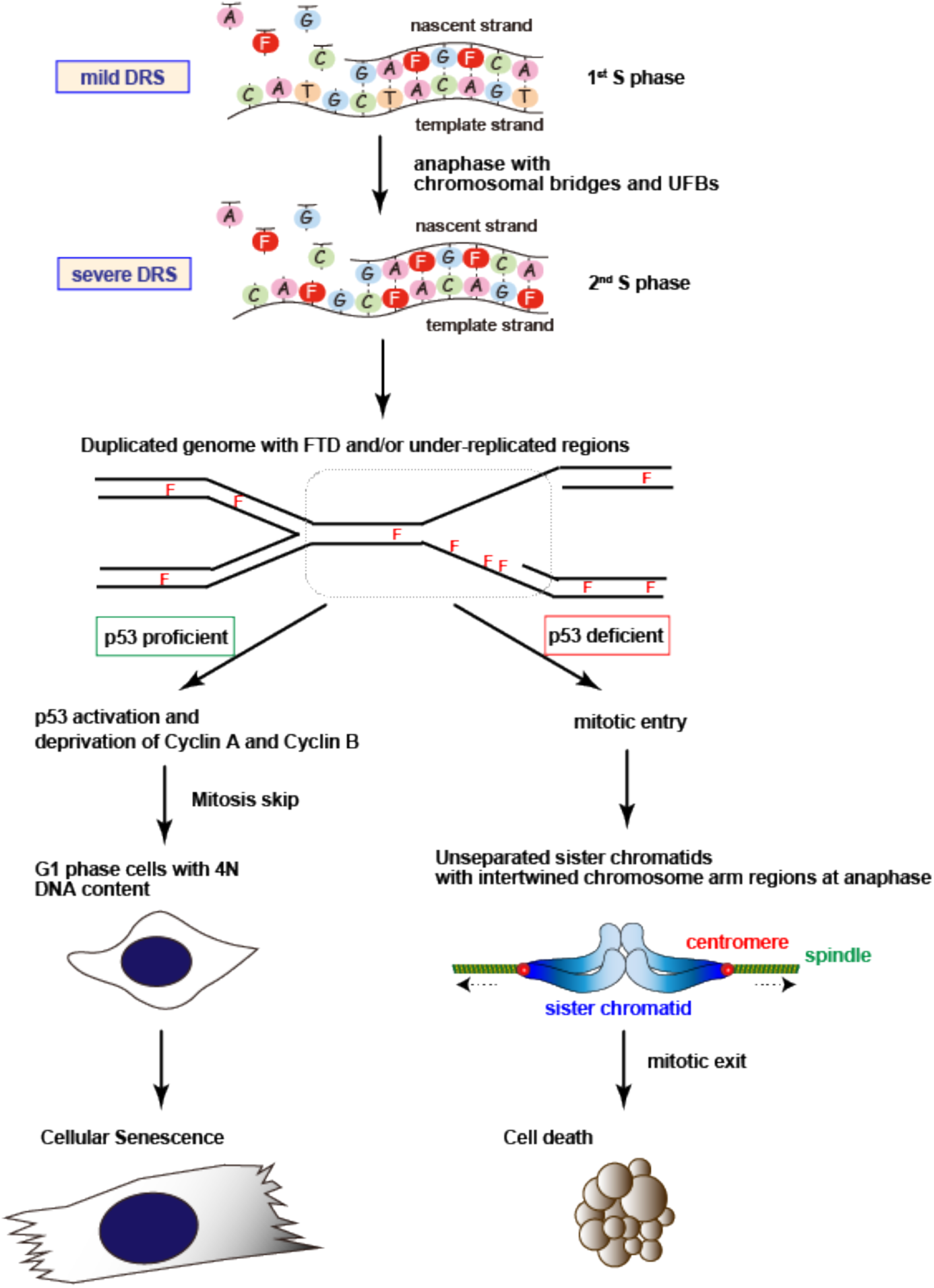
Schematic model of the effect of FTD and cell fate decision in tumor cells. See text for details.

Biochemical analysis of the *in vitro* reconstituted DNA replication process using human Polδ and Polε revealed the unique properties of FTD as a nucleoside analog. First, as a component of the dNTP pool, FTD-TP can replace dTTP during DNA synthesis at the replication fork; however, the incorporation of FTD-TP was not efficient. Another intriguing property of FTD-TP is that FTD incorporation did not terminate DNA polymerization, allowing its continuous incorporation into DNA to produce FTD-containing DNA strands. By contrast, other chemotherapeutic agents, such as gemcitabine and cytarabine, terminate DNA polymerization in the vicinity of their incorporation (16) and are incorporated into DNA to a lower extent than FTD (40). Second, as a component of the template DNA strand, FTD constitutes a continuous obstacle to DNA polymerization. This property may explain the severe extension of the second S/G2 phase after FTD addition, because DNA synthesis at the second S phase would have to proceed with the FTD-containing template DNA strand in the presence of FTD-TP in the dNTP pool. Third, FTD incorporated into the DNA of tumor cells is retained for a prolonged period (41). This property would cause persistent disturbance of 19 the replication process and tumor cytotoxicity, which may underlie the sustained growth suppressive effect and prolonged survival observed in a xenograft mouse model exposed to limited courses of FTD/TPI (40). The dual perturbation of DNA replication caused by the inefficient incorporation of FTD during DNA synthesis and the persistent roadblock, which represents FTD incorporated into template DNA strand, for DNA polymerization are probably key properties underlying the success of this chemotherapeutic drug.

Because FTD is a thymidine analog, inefficient DNA replication occurs preferentially at AT-rich genomic loci, *e.g*. CFSs (42). FTD would enhance DRS at these fragile sites and exacerbate DNA lesions resulting from the generation of UR-DNA. However, our previous study did not detect an increase in DNA strand breaks in FTD-treated HCT-116 cells (21). DNA lesions such as UR-DNA with FTD incorporation at CFSs could be protected by proteins that accumulate at these lesions, such as FancD2 and the RPA complex, as suggested by the formation of discrete FancD2 and RPA32 nuclear foci (Fig. 3C), and result in the activation of the p53-p21 pathway to avoid catastrophic collapse of chromosomes (14). On the other hand, in p53 knock-out cells, such DNA lesions were probably converted to detrimental DNA strand breaks, as evidenced by the detection of γH2AX (Fig. 4E), through aberrant mitosis progression and unseparated sister chromatids probably caused by interlinks between UR-DNA located at chromosomal arms (Fig. 6).

In this study, cellular analysis revealed that FTD suppresses the growth of tumor cells irrespective of p53 status. p53-dependence diverged at the second G2/M phase transition, resulting the redirection of cell fate toward either cellular senescence or apoptotic cell death. p53-mediated senescence, however, impairs the apoptotic response to chemotherapy, and the senescent tumor cells show persistent mitogenic potential, which causes relapse (43). Recent studies indicate that establishment of senescence may reprogram tumor cells into a latent stem-like state, resulting in tumor cells that escape senescence having a more aggressive 20 phenotype (44,45). Because FTD is a component of the chemotherapeutic drug FTD/TPI (18-20), it is critical to determine whether FTD-induced senescent tumor cells also acquire stemness and a persistent mitogenic potential. If that is the case, a strategy for evading FTD-induced senescence or for directing tumor cell fate toward apoptotic death should be given serious consideration.

FTD/TPI shows efficacy in the treatment of colorectal cancer patients who are refractory or intolerant of 5-FU-based therapy (17-19). At the cellular level, acquired resistance to 5-FU does not confer FTD resistance and *vice versa* (22,46,47), possibly reflecting the distinct mechanism of action of each drug. The unique and distinct property of FTD demonstrated in this study may expand the clinical application of FTD-based therapy. Additionally, gain-of-function mutations or heterozygous mutations are frequently found in the *TP53* gene locus of tumors obtained from patients (48). Therefore, investigating whether the expression of mutant p53 affects the cellular response and tumor cell fate decision induced by FTD is an important issue.

## ACKNOWLEDGEMENTS

We would like to thank Ms. Masako Kosugi, Tomomi Takada, Naoko Katakura, and Atsuko Yamaguchi for their expert technical assistance; Dr. Yoko Katsuki for the technical advice; and Dr. Mamoru Kiniwa and Dr. Minoru Takata for critical reading of the manuscript. We also appreciate the technical assistance from the Research Support Center, Research Center for Human Disease Modeling, Kyushu University Graduate School of Medical Sciences.

## AUTHOR CONTRIBUTIONS

M.I. and H.K. conceived the project. Y.K., M.I., R.F., T.T., T.M-I., D.M., and H.K. developed the methodologies. Y.K., M.I., S.N., T.W., and H.K. performed and analyzed the cellular response experiments; R.F. and T.T. performed and analyzed *in vitro* DNA replication experiments; T.M-I. and D.M. performed and analyzed metabolomic experiments; H.S., and E.O. analyzed the data and contributed to the discussion from the clinical point-of-view. H.K. wrote the manuscript and Y.K., M.I., T.W., and H.K. edited the manuscript. Y.M., and H.K. supervised this study.

## SUPPLEMENTARY DATA

Fig. S1. Detection of nucleoside analogs by immunofluorescence staining.

Fig. S2. Experimental design of the *in vitro* DNA synthesis assay.

Fig. S3. *In vitro* human DNA replication assay related to Fig. 1G.

Fig. S4. Data related to Fig. 2.

Fig. S5. Gene targeting of the *TP53* gene by CRISPR/Cas9.

Fig. S6. Data related to Fig. 4A.

Fig. S7. Cellular responses of A549 and DLD1 cells.

Fig. S8. Immunofluorescence images. Data related to Fig. 6C.

