## Supplemental Table 1 for "DNA replication stress induced by trifluridine determines tumor cell fate according to p53 status"

Table S1. The list of antibodies specific to antigens

| Experiment | Antigen | Supplier | Cat # | Dilution |
|---|---|---|---|---|
| DNA fiber | BrdU (B44) | BD Bioscience | 347580 | 1:25 |
| | BrdU (BU1/75) | Abcam | ab6326 | 1:400 |
| Western blot | FancD2 | Novus Biologicals | NB 100-182 | 1:1,000 |
| | PARP | Cell Signaling Technology | 9542 | 1:1,000 |
| | cleaved PARP | Cell Signaling Technology | 5625 | 1:1,000 |
| | Chk1 | Santa Cruz Biotechnology | sc-8408 | 1:1,000 |
| | phospho Ser345 Chk1 | Cell Signaling Technology | 2348 | 1:500 |
| | Cyclin A | Merk Millipore | 05-374 | 1:1,000 |
| | Cyclin B1 | Merk Millipore | 05-373 | 1:1,000 |
| | Cyclin D1 | Santa Cruz Biotechnology | sc-20044 | 1:500 |
| | p53 | DAKO | M7001 | 1:1,000 |
| | p21 | Santa Cruz Biotechnology | sc-397 | 1:1,000 |
| | Caspase 3 | Cell Signaling Technology | 9662 | 1:1,000 |
| | cleaved Caspase 3 | Cell Signaling Technology | 9664 | 1:1,000 |
| | histone H3 | Abcam | 1791 | 1:2,000 |
| | γH2AX | Merk Millipore | 05-636 | 1:1,000 |
| | PCNA | Epitomics | 2714-1 | 1:10,000 |
| | β-actin | Sigma | A5316 | 1:5,000 |
| Immunofluorescence | PICH | Merk Millipore | 04-1540 | 1:50 |
| | α-tubulin | Sigma | T6199 | 1:2,000 |
| | centromere (ACA) | Immunovision | HCT-0100 | 1:10,000 |
| | Cyclin B1 | Merk Millipore | 05-373 | 1:500 |
| | FancD2 | Novus | NB 100-182 | 1:500 |
| | RPA32 | Abcam | ab2175 | 1:200 |
