## Supplemental Figures 1-8 for "DNA replication stress induced by trifluridine determines tumor cell fate according to p53 status"

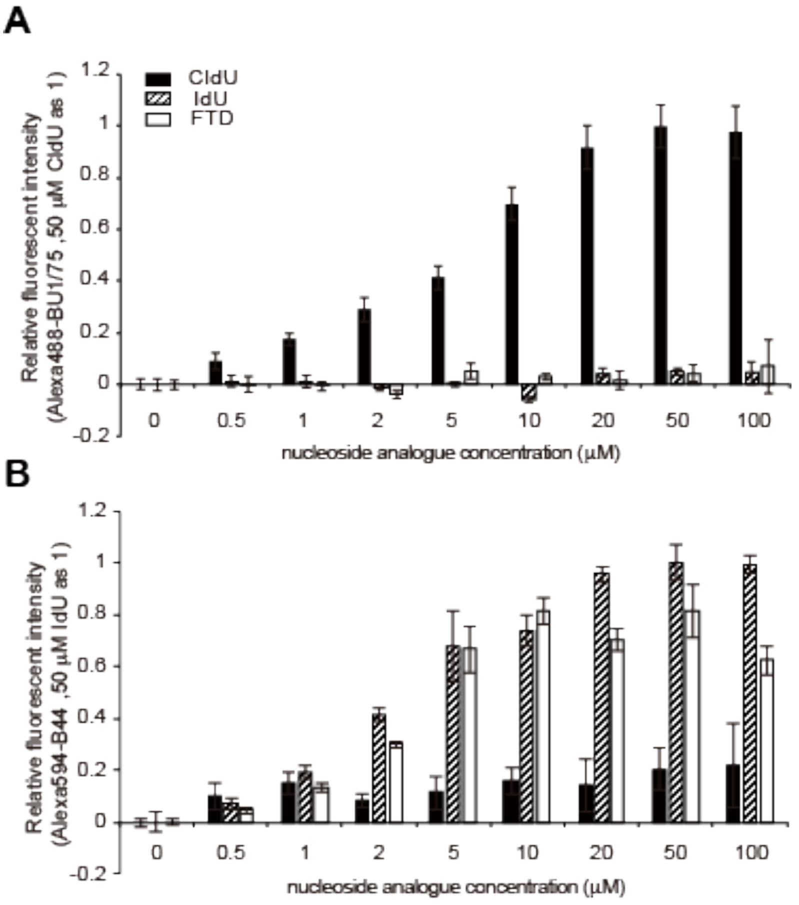


**Fig. S1**

**Fig. S1.** Detection of nucleoside analogs by immunofluorescence staining. (**A**) Detection with the anti-BrdU antibody BU1/75. HCT-116 cells were cultured in the presence of CldU, IdU or FTD at the indicated concentrations for 1 h and immunostained with the anti-BrdU antibody BU1/75. Relative fluorescence intensity was calculated considering the average amount of 50 μM CldU as 1. (**B**) Detection with the anti-BrdU antibody B44. Relative fluorescence intensity was calculated considering the average amount of 50 μM IdU as 1.

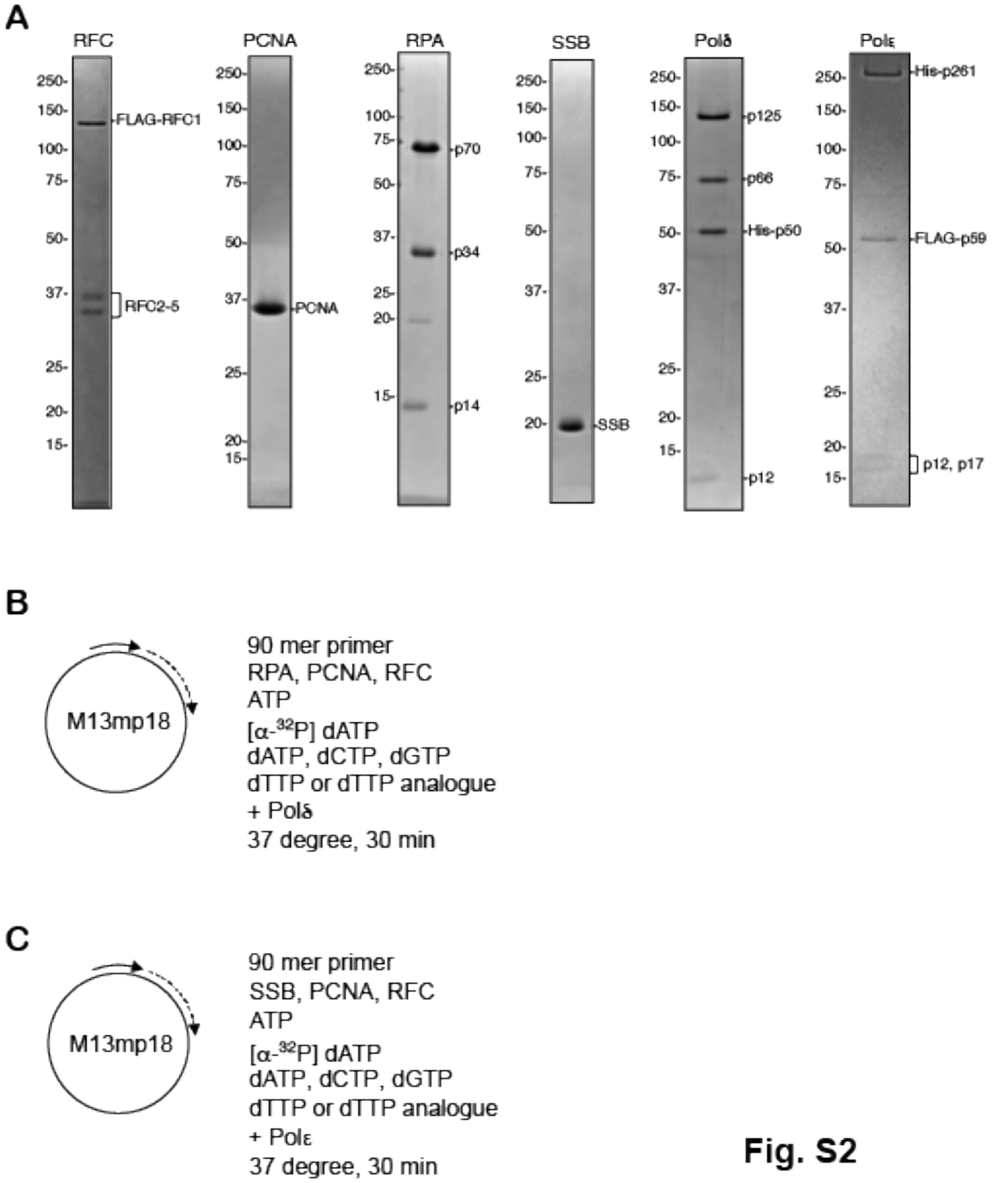


Fig. S2

**Fig. S2.** Experimental design of the *in vitro* DNA synthesis assay. (**A**) Purified proteins. Coomassie blue staining of each purified protein or protein complex is shown. The protein purification methods are described in Materials and methods. (**B**) Schematic of the *in vitro* DNA synthesis assay using Polδ. **C.** Schematic of the *in vitro* DNA synthesis assay using Polε.

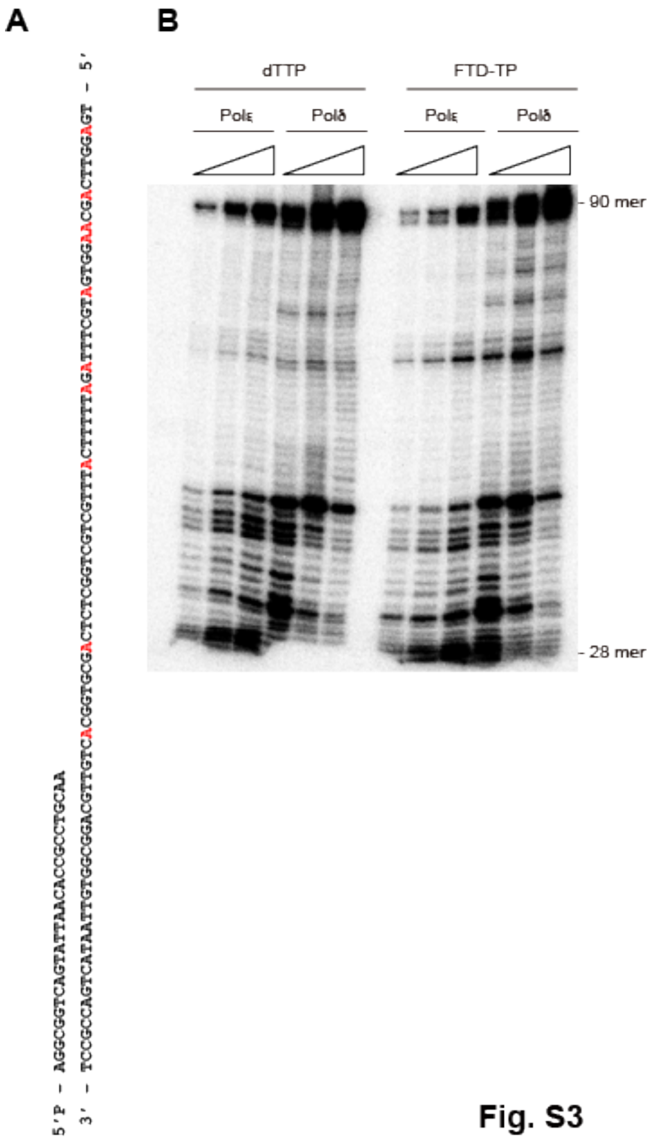


Fig. S3

**Fig. S3.** *In vitro* DNA synthesis assay related to Fig. 1G. (**A**) Sequences of 5′-radiolabeled primer and template 90-mer oligonucleotides of the *in vitro* DNA synthesis analysis. The A sequences in the template oligonucleotide are marked by red. (**B**) DNA synthesis from radiolabeled primers of the 90-mer oligonucleotide template. The DNA synthesis reactions catalyzed by Polδ and Polε were performed in the presence of dTTP or FTD-TP.

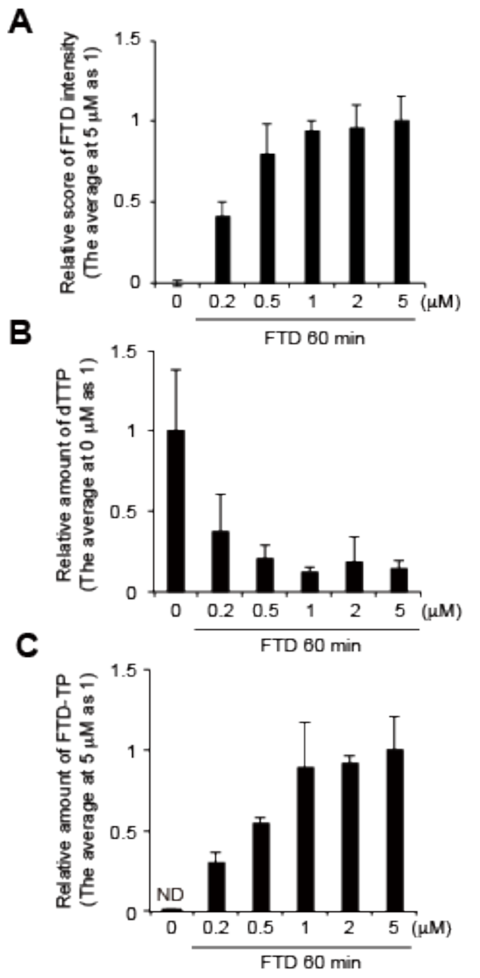


**Fig. S4**

**Fig. S4.** Data related to Fig. 2. (**A**) FTD incorporation into DNA. The relative fluorescence intensity was calculated considering the average amount of 5 μM FTD as 1. (**B,C**) Relative amount of cellular dTTP (**B**) and FTD-TP (**C**). The relative scores were calculated considering the average amount of 0 μM FTD (**B**) or 5 μM FTD (**C**) as 1.

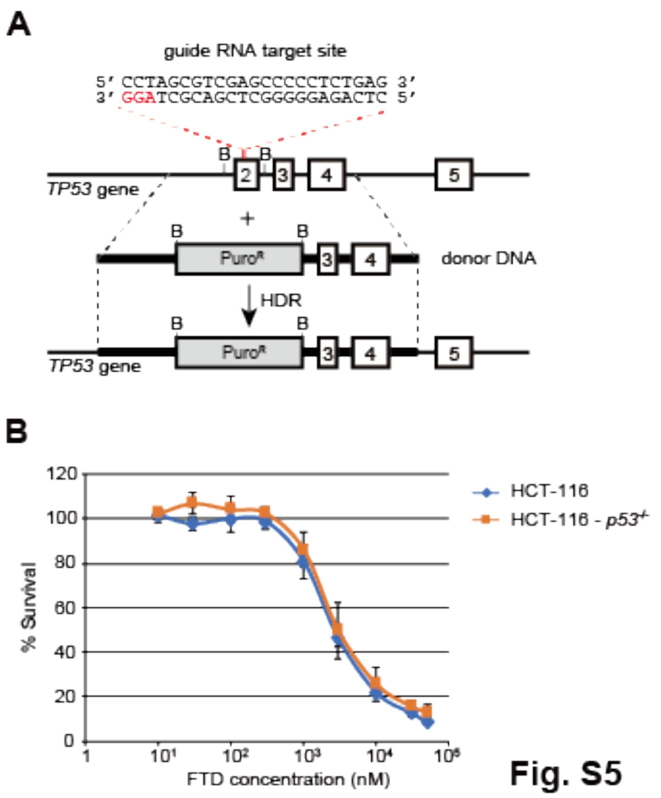


Fig. S5

**Fig. S5.** Gene targeting of the *TP53* gene by CRISPR/Cas9. (**A**) Schematic design of CRISPR/Cas9-mediated gene targeting. The position and sequence of the guide RNA, donor DNA, and the targeted allele are shown. B indicates the *Bam* HI site. (**B**) Viability of HCT-116 and HCT-116 $p53^{-/-}$ cells in the presence of various concentrations of FTD on day 3.

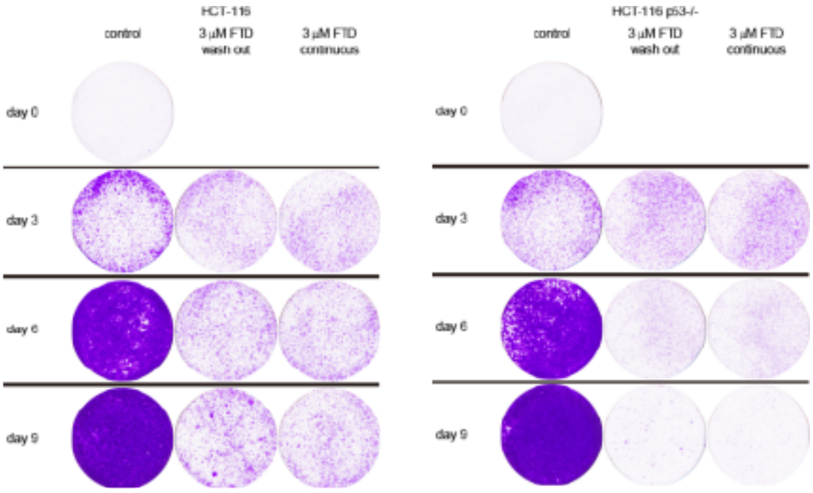


**Fig. S6**

**Fig. S6.** Data related to Fig. 3A. Raw imgaes of crystal violet staining of HCT-116 or HCT-116 *p53*$^{-/-}$ cells cultured in the presence of FTD.

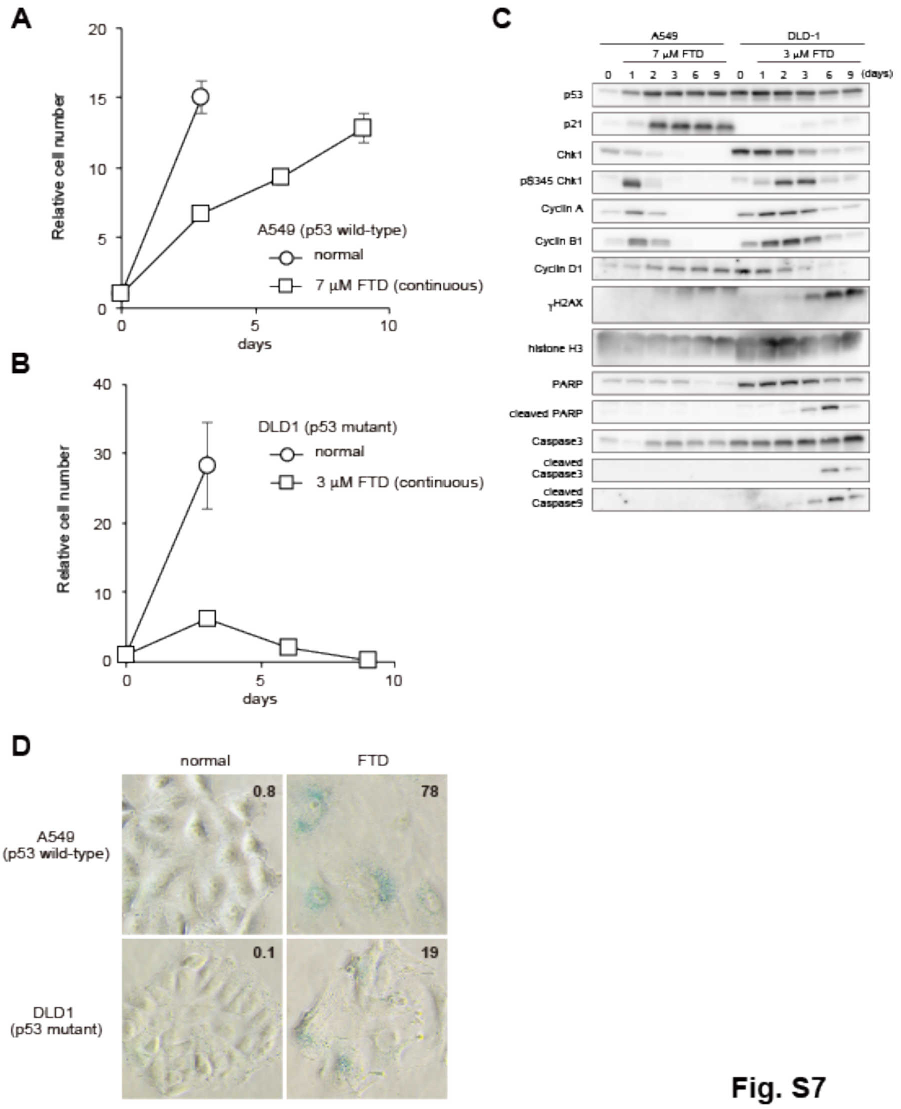


**Fig. S7**

**Fig. S7.** Cellular responses of A549 and DLD1 cells. Proliferation rate (A549: **A,** DLD1: **B**), DNA damage response (**C**) and senescence associated β-galactosidase (SA-β-gal) activity on day 3 (**D**) in A549 and DLD1 cells cultured in the presence of the $IC_{50}$ concentration of FTD.

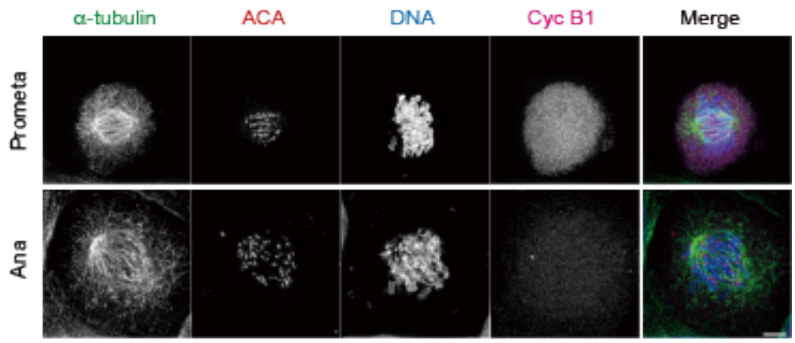


**Fig. S8**

**Fig. S8.** Immunofluorescence images. Data related to Fig. 5C. Fluorescence immunostaining of α-tubulin, centromere (ACA), cyclin B1 and DNA in proliferating HCT-116 cells at prometaphase (Prometa) and anaphase (Ana).
